# Engineering Yeast to Improve Heterologous Abscisic Acid Production

**DOI:** 10.1101/2023.06.07.544016

**Authors:** Maximilian Otto, Michael Gossing, Florian David, Verena Siewers

## Abstract

Abscisic acid (ABA) is a high-value product with agricultural, medical and nutritional applications. We previously constructed an ABA cell factory by expressing the ABA metabolic pathway from *Botrytis cinerea* in the biotechnological workhorse *Saccharomyces cerevisiae*.

In this study, we aimed to improve ABA production and explored various rational engineering targets mostly focusing on increasing the activity of two rate-limiting cytochrome P450 monooxygenases of the ABA pathway, BcABA1 and BcABA2. We evaluated the effects of overexpression and knock-down of cell membrane transporters, expression of heterologous cytochrome b5, overexpression of a rate-limiting heme biosynthesis gene and overexpression or knock-out of genes involved in ER membrane homeostasis. One of the genes involved in ER membrane homeostasis, *PAH1*, was identified as the most promising engineering target. Knock-out of *PAH1* improved ABA titers, but also caused a sever growth defect. By replacing the *PAH1* promoter with a weak minimal promoter, it was possible to mediate the growth defect while still improving ABA production.

In this report we were able to improve the ABA cell factory and furthermore provide valuable insights for future studies aiming to engineer cytochrome P450 monooxygenases.

**One-sentence summary:** In this study we explored various strategies to improve heterologous abscisic acid production in *Saccharomyces cerevisiae* and identified fine-tuning of the *PAH1* gene as a promising engineering strategy.

## Introduction

The isoprenoid abscisic acid (ABA) has become a molecule-of-interest for a multitude of applications in agriculture and medicine as well as nutrition. Its central role in plant physiology has long been known and is still being investigated (Finkelstein 2013; Chen *et al*. 2020). Agricultural applications of ABA range from alleviating various abiotic stresses to regulating seed dormancy, germination and fruit ripening (Sah, Reddy and Li 2016; Vishwakarma *et al*. 2017; Gupta *et al*. 2022). ABA acts as a signalling molecule in many other organisms besides plants (Olds, Glennon and Luckhart 2018). In recent years, ABA was shown to have various biological activities in mammals, making it a promising drug candidate. Pharmacological activity against various inflammatory diseases, pathogen-mediated infections, type-2-diabetes and metabolic syndrome (a complex condition with various comorbidities) has been reported (Sakthivel *et al*. 2016; Lievens *et al*. 2017; Kim *et al*. 2020). ABA was furthermore identified as a bitter receptor blocker with potential applications in nutritional products (Pydi *et al*. 2015).

A sustainable and inexpensive product source will be required to utilise ABA in these applications. The plant pathogenic fungus *Botrytis cinerea* is a natural producer of ABA and has been used as a biotechnological production host (Shi *et al*. 2016). In *B. cinerea*, ABA is produced via the mevalonate pathway from farnesyl-pyrophosphate (FPP) (Hirai *et al*. 2014). FPP is cyclised by the enzyme BcABA3 and subsequently oxidised at multiple positions by BcABA1, BcABA2 and BcABA4 to form ABA (Figure 1) (Inomata *et al*. 2004; Siewers, Smedsgaard and Tudzynski 2004; Siewers *et al*. 2006; Takino *et al*. 2019). However, the lack of genetic tools for *B. cinerea*, as well as its hyphal morphology and comparatively slow growth make the fungus a challenging cell factory host.

**Figure 1:**
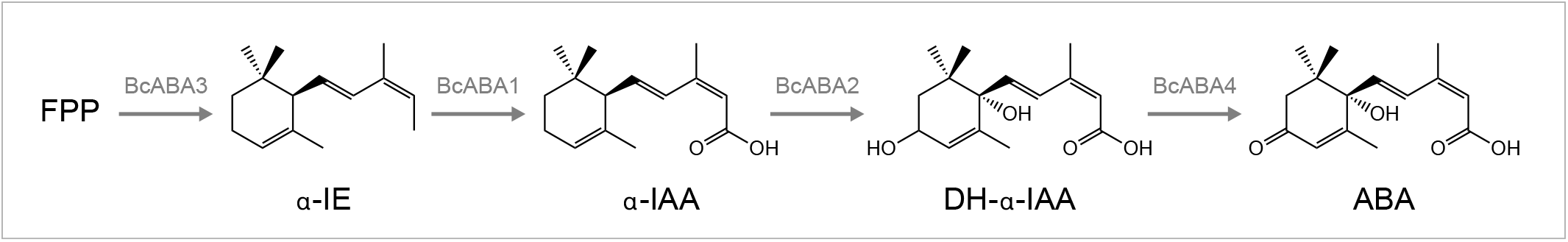
ABA pathway in *Botrytis cinerea* (Takino et al. 2019). The four pathway genes *bcaba1, bcaba2, bcaba3* and *bcaba4*, as well as the *B. cinerea* cytochrome P450 reductase encoding gene *bccpr1* were expressed in yeast to enable heterologous ABA production from the native precursor FPP (Otto et al. 2019). Abbreviations: FPP = farnesyl pyrophosphate, α-IE = α-ionylideneethane, α-IAA = α-ionylideneacetic acid, DH-α-IE = 1’,4’-trans-dihydroxy-α-ionylideneacetic acid, ABA = abscisic acid

*Saccharomyces cerevisiae* has long been a workhorse for biotechnological applications. Its capabilities for high-level production of sesquiterpenes or sesquiterpenoids have been demonstrated previously for the biofuel farnesene (Meadows *et al*. 2016) and the anti-malarial drug artemisinic acid (Paddon *et al*. 2013). In a previous study, we established an *S. cerevisiae* ABA cell factory by expressing the ABA pathway enzymes BcABA1, BcABA2, BcABA3 and BcABA4 as well as the *B. cinerea* cytochrome P450 reductase BcCPR1 (Otto *et al*. 2019). We demonstrated that the two cytochrome P450 monooxygenases (CYPs, putatively of class II) BcABA1 and BcABA2 are limiting production in the current strain (Otto *et al*. 2019).

CYPs are a large superfamily of enzymes that catalyse oxidation reactions using molecular oxygen and contain heme as a co-factor (Denisov *et al*. 2005). They are ubiquitous in plant and microbial biosynthetic pathways and have become a major engineering target in biotechnology (Podust and Sherman 2012; Jiang *et al*. 2021). Class II CYP enzymes, the most common class in eukaryotes, require cytochrome P450 reductases (CPRs) as co-enzymes for the transfer of electrons from NADPH (Werck-Reichhart and Feyereisen 2000). It was shown that co-expression of cytochrome b5 (CYB5) and its cognate reductase (CBR) can have a positive impact on CYP activities, most likely by acting as additional electron donors (Schenkman and Jansson 2003; Zhang, Im and Waskell 2007; Paddon *et al*. 2013). CYPs, CPRs, CYB5s and CBRs are anchored in the ER membrane.

Expansion of the ER membrane structure, also referred to as ER proliferation, has developed into a common engineering strategy for membrane-associated enzymes like CYPs and their co-enzymes (Jiang *et al*. 2021). It is presumed that available membrane space is often limiting enzyme abundance in the cell. Various native target genes have been investigated that can cause ER proliferation, some of which lead to multi-fold titer increases

(Arendt *et al*. 2017; Kim *et al*. 2019). However, the effect of individual target genes appears to be strain or product specific and studies comparing multiple targets are rare.

Besides ER membrane space, heme availability can limit production in some CYP-overexpressing cell factories (Michener, Nielsen and Smolke 2012; Park and Choi 2020). Native heme biosynthesis in yeast provides an additional engineering target to improve CYP activity.

When produced in yeast, ABA is predominantly found in the cell culture supernatant (Otto *et al*. 2019). As a weak organic acid with a *pK*_a_ of 4.75 (Slovik, Baier and Hartung 1992), ABA is mostly present as the conjugate base in the cytosol of plants, with the membrane permeability of the ion being much lower than its protonated form (Heilmann, Hartung and Gimmler 1980; Kaiser and Hartung 1981). In yeast, ABA is presumably being exported by an unspecific yeast transporter. Complex regulatory networks mediate resistance to weak organic acids in *S. cerevisiae* with a multitude of transporters being involved in this process (Mira, Teixeira and Sá-Correia 2010). Upregulation of native transporters have been investigated as engineering targets for various cell factories with the goal of mediating product-related cell stress, such as increased turgor pressure or oxidative stress (Mira, Teixeira and Sá-Correia 2010). Nonetheless, downregulation of transporters could also be a valid engineering target by preventing the export of pathway intermediates, increasing their local concentration in the cytosol. Finding transporters involved in the import or export of a given heterologous compound is challenging. An alternative approach is to overexpress or knock-out of global regulators involved in transporter expression. *PDR1* and *YRR1* are two such global transcriptional regulators, involved in the pleiotropic drug resistance (PDR) signalling network (Kolaczkowska and Goffeau 1999; Le Crom *et al*. 2002). Interfering with the PDR network could provide valuable insight for ABA-producing yeast strains and other cell factories producing weak organic acids.

Our previous proof-of-concept study demonstrated the feasibility of an *S. cerevisiae* cell factory, but ABA titres remained low. Here, we aimed to rationally engineer the native yeast metabolism to improve ABA production by focusing on CYP activity as well as unspecific yeast transporters.

## Materials and Methods

### PCRs and plasmid construction

For PCR reactions PrimeSTAR HS DNA Polymerase (Clontech), Phusion High-Fidelity DNA polymerase (Thermo Fisher Scientific) or SapphireAmp (Takara Bio) were used and the manufacturer ‘s instructions were followed. Primer sequences can be found in Supplementary Table S1 and specific conditions of standard PCR reactions are listed Supplementary Table S2. For plasmid and PCR product purification, GeneJet Purification Kits (Thermo Fisher Scientific) were used. Sanger DNA sequencing was performed by Eurofins Genomics. Primers and short oligos were ordered from Eurofins Genomics or Integrated DNA Technologies.

Plasmids used in this study ‘s experiments are listed in Table 2. The Golden Gate-based MoClo workflow (Engler, Kandzia and Marillonnet 2008; Lee *et al*. 2015; Otto *et al*. 2021) was followed to construct pX3-bcaba1+2 and pXII2-bccyb5+cbr1. All MoClo assemblies are listed in Supplementary Table S3. First, primers pairs 306/357 and 307/310 were used to remove a BsaI site from *bcaba1* and to attach MoClo type-3 compatible overhangs via PCR. The resulting PCR products were fused together via a 2-step PCR reaction. In the first reaction, the PCR products were mixed in a 1:1 ratio (≈100 ng) with PrimeStar PCR master mix but without primers (55 °C T_ann_, 1:15 min t_elo_, 15 cycles). In the second reaction, 2 µL of the first reaction was used as template with the primer pair 306/307 (55 °C T_ann_, 1:40 min t_elo_, 35 cycles). The PCR product of the second reaction was purified and sequence verified. Primer pairs 308/309 were used to attach type-3 MoClo overhangs to *bcaba2*.

DNA sequences lacking BsaI, BsmbI and NotI sites for *bccyb5* (UniProt identifier A0A384K1M2) and *bccbr1* (UniProt identifier A0A384JNH0) were codon-optimised for yeast and provided by Genscript Biotech Corp. Primer pairs 292/293 and 294/295 were used to attach MoClo type-3 overhangs to *bccyb5* and *bccbr1*, respectively.

The MoClo compatible *bcaba1, bcaba2, bccyb5* and *bccbr1* fragments were inserted into the MoClo entry vector pYTK001 (Lee *et al*. 2015) (Supp. Table S3). Subsequently, level-1 MoClo plasmids were assembled for each gene. The level-1 plasmids were combined with the backbones pMC-X3 and pMC-XII2 (Otto *et al*. 2021) to from pX3-bcaba1+2 and pXII2-bccyb5+cbr1, respectively.

### Microorganisms and media

Plasmids were amplified using NEB® 5-alpha Competent *E. coli* cells (New England Biolabs). *S. cerevisiae* strains are listed in Table 1 and a pedigree chart is shown in Figure 2. *E. coli* was cultivated at 37 °C in liquid Luria-Bertani (LB) media or on LB agar plates with appropriate antibiotics.

**Table 1:**
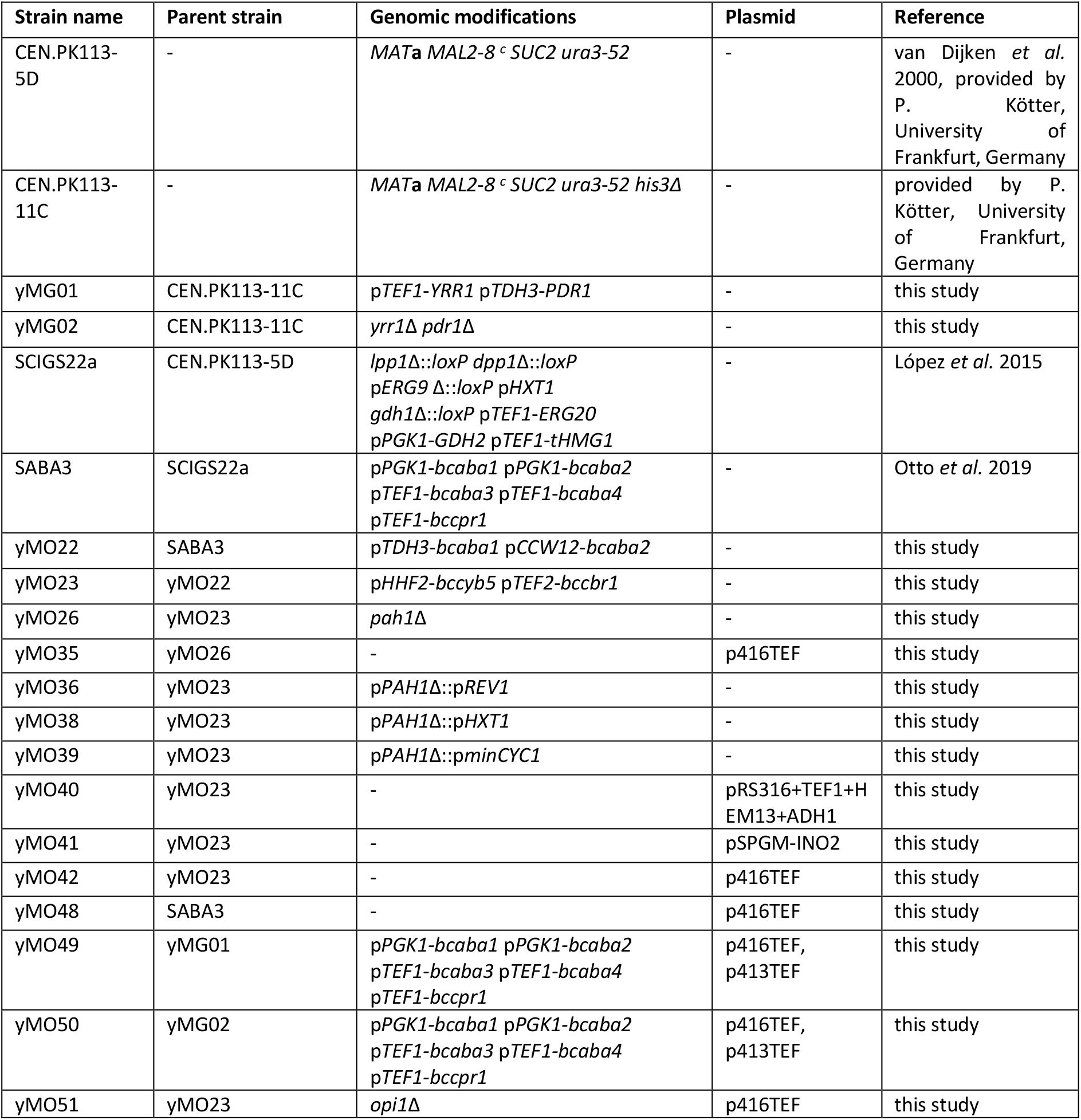
Strains used and constructed in this study

**Table 2:** Plasmids used in this study

| Plasmid name | Use | Backbone | Expression cassette | Reference |
| --- | --- | --- | --- | --- |
| p416TEF | Episomal expression | - | empty | Mumberg, Müller and Funk 1995 |
| p413TEF | Episomal expression | - | empty | Mumberg, Müller and Funk 1995 |
| pRS316+TEF1+HEM13+ADH1 | Episomal expression | pRS316 | pTEF1-HEM13 | Ishchuk *et al.* 2022 |
| pSPGM-INO2 | Episomal expression | pSP-GM1 | pTEF1-INO2 | Chen *et al.* 2022 |
| pCfB2904-ABA1-CPR | Genomic integration XI-3 | pCfB2904 | pPGK1-bcaba1 pTEF1-bccpr1 | Otto *et al.* 2019 |
| pCfB2909-ABA2-ABA4 | Genomic integration in XII-5 | pCfB2909 | pPGK1-bcaba2 pTEF1-bcaba4 | Otto *et al.* 2019 |
| pCfB3035-ABA3 | Genomic integration in X-4 | pCfB3035 | pTEF1-bcaba3 | Otto *et al.* 2019 |
| pX3-bcaba1+2 | Genomic integration in X-3 | pMC-X3 | pTDH3-bcaba1 pCCW12-bcaba2 | This study |
| pXII2-bccyb5+cbr1 | Genomic integration in XII-2 | pMC-XII2 | pHHF2-bccyb5 pTEF2-bccbr1 | This study |

**Figure 2:**
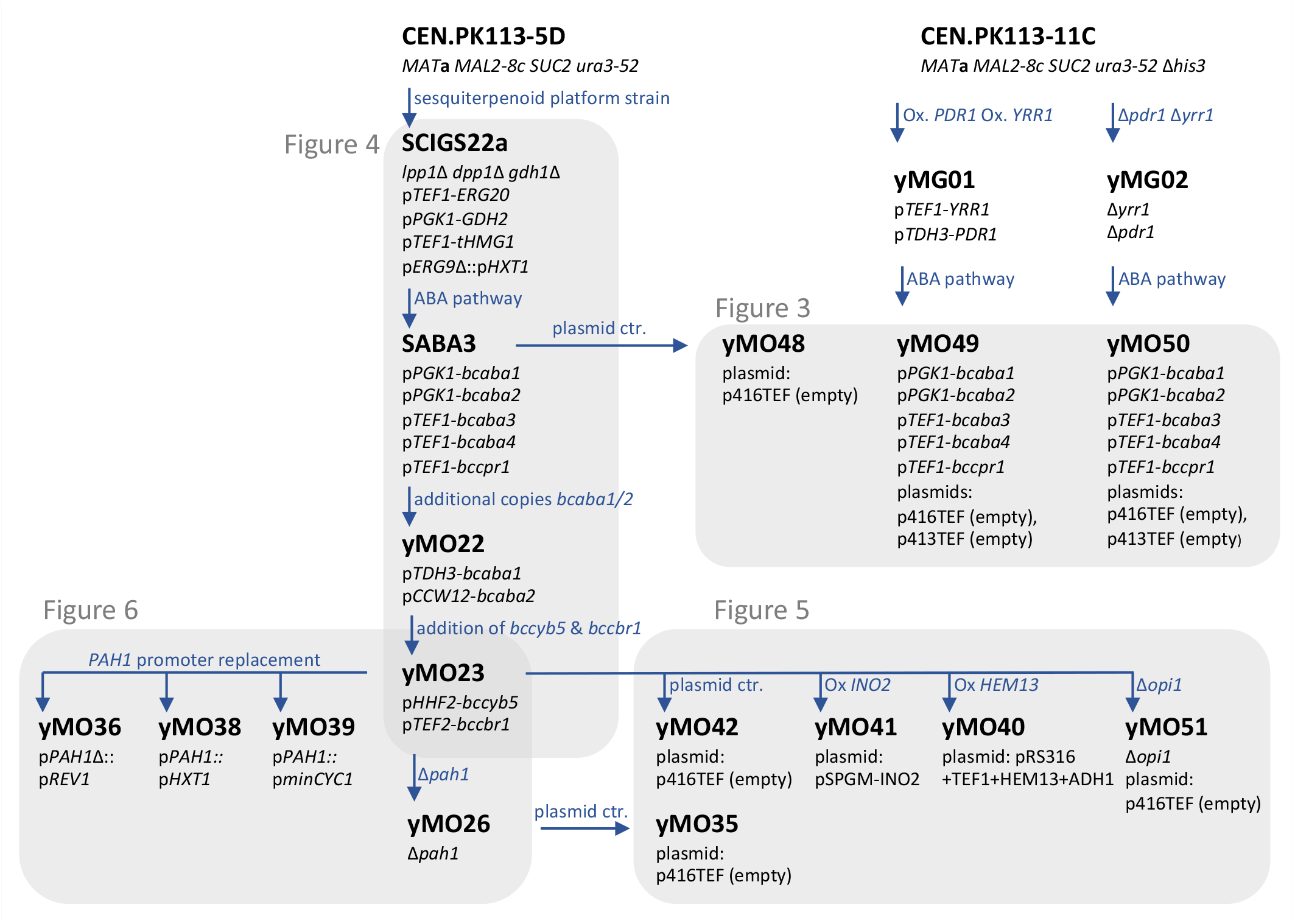
Strain pedigree tree of parent strains and strains constructed in this study. Grey boxes indicate strains that were compared in Figures 3-6. For more detailed strain information see Table 1 in Materials and Methods section. Ox = overexpression

Yeast was cultivated at 30 °C in liquid yeast extract peptone dextrose (YPD) media or mineral media (adapted from Verduyn *et al*. 1992) shaking at 220 rpm. For cultivation on agar plates YPD media or synthetic defined (SD) media was used. 20 g/L glucose was used for all yeast media. Detailed media compositions can be found in Supplementary Table S4.

### Strain construction

Yeast strains were transformed according to the protocol from Gietz and Woods (2006). The plasmids pX3-bcaba1+2 and pXII2-bccyb5+cbr1 were used for genomic integrations. The procedure was followed as described in Jessop-Fabre *et al*. (2016), using the Cas9-encoding plasmid pCfB2312 and the gRNA-helper plasmids pCfB3041 and pCfB3051.

*PAH1* and *OPI1* were knocked out following the protocol of Mans *et al*. (2015), using the plasmid pMEL10 for gRNA expression and the Yeastriction tool (http://yeastriction.tnw.tudelft.nl/). The sequence of the used gRNA fragments and repair oligos can be found in Supplementary Table S5. Primers pairs 366/367 and 1260/1261 (Supp. Table S1 and Supp. Table S2) were used to validate the Δ*pah1* and Δ*opi1* genotype, respectively.

For the *PAH1* promoter replacement, the gRNA was designed using the Benchling CRISPR Guide tool (Supp. Table S5) and its coding sequence inserted into pMEL10. For construction of the linear promoter replacement cassettes, primers pairs 417/418, 422/423, 424/425 were used. The primers were designed to contain homologous regions for replacing 771 base pairs upstream of the *PAH1* start codon. The resulting PCR products were used as repair fragments in a transformation according to Mans *et al*. (2015). Primers 426/427 were used to validate the p*PAH1* replacement. The colony PCR product of strain yMO36 was sequenced since the wild-type p*PAH1* band and p*REV1*-*PAH1* band are hard to distinguish by size.

### Cultivation for OD_600_ and ABA analysis

For the experiments displayed in Figure 3-6, single colonies were picked from agar plates for precultures (1.5 mL mineral media) in 14 mL round-bottom cultivation tube (Greiner Bio-One). Precultures were grown for 48 or 72 h (for slow growing Δ*pah1* strains) and main cultures (2.5 mL mineral media) were inoculated at OD_600_ 0.1 in 24-deepwell microplates (square wells, pyramid-bottom, EnzyScreen). Cultures were grown for 48 h (Figure 3, Figure 4) or 60 h (Figure 5, Figure 6), OD_600_ was measured, cultures were centrifuged (5 min, 1500 xg) and 1 mL supernatant was transferred to 2-mL Eppendorf tubes.

**Figure 3:**
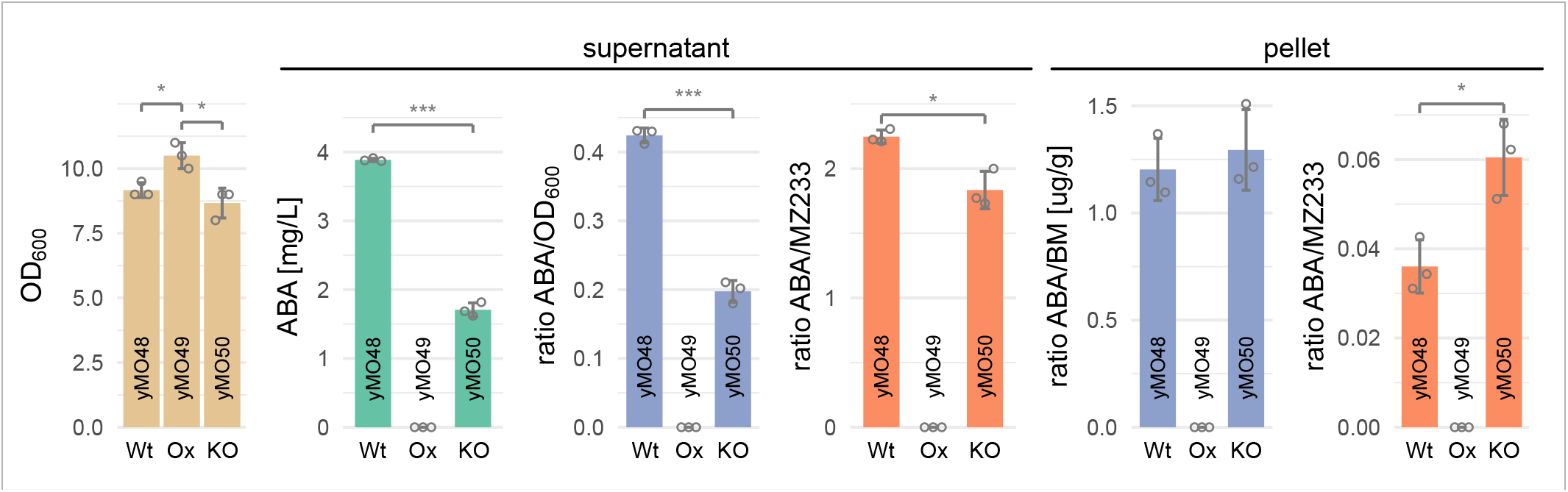
Effect of native transporter overexpression or knockdown in ABA-producing strains. Genes encoding the transcription factors Pdr1 and Yrr1 were either overexpressed (yMO49) or knocked out (yMO50) and compared to a control strain with wild-type *PDR1* and *YRR1* expression (yMO48). OD600 (beige), ABA titer (green), ABA titer normalized to OD600 or biomass (blue), and ion count ratio of ABA to MZ233 (orange) are shown after 48 h of cultivation in mineral media (24-deepwell microplates). Supernatant and cell pellet were analysed separately. Grey circles show the individual data points used to calculate the mean and standard deviation. Only traces of ABA (<0.05 mg/L) and MZ233 were detected for yMO49. Significance (Student ‘s t-test, two-sided) is displayed as “* “ and “*** ‘ ‘, indicating p<0.05 and p<0.001 respectively. Abbreviations: Wt = wild-type, Ox = overexpression, KO = knockout

**Figure 4:**
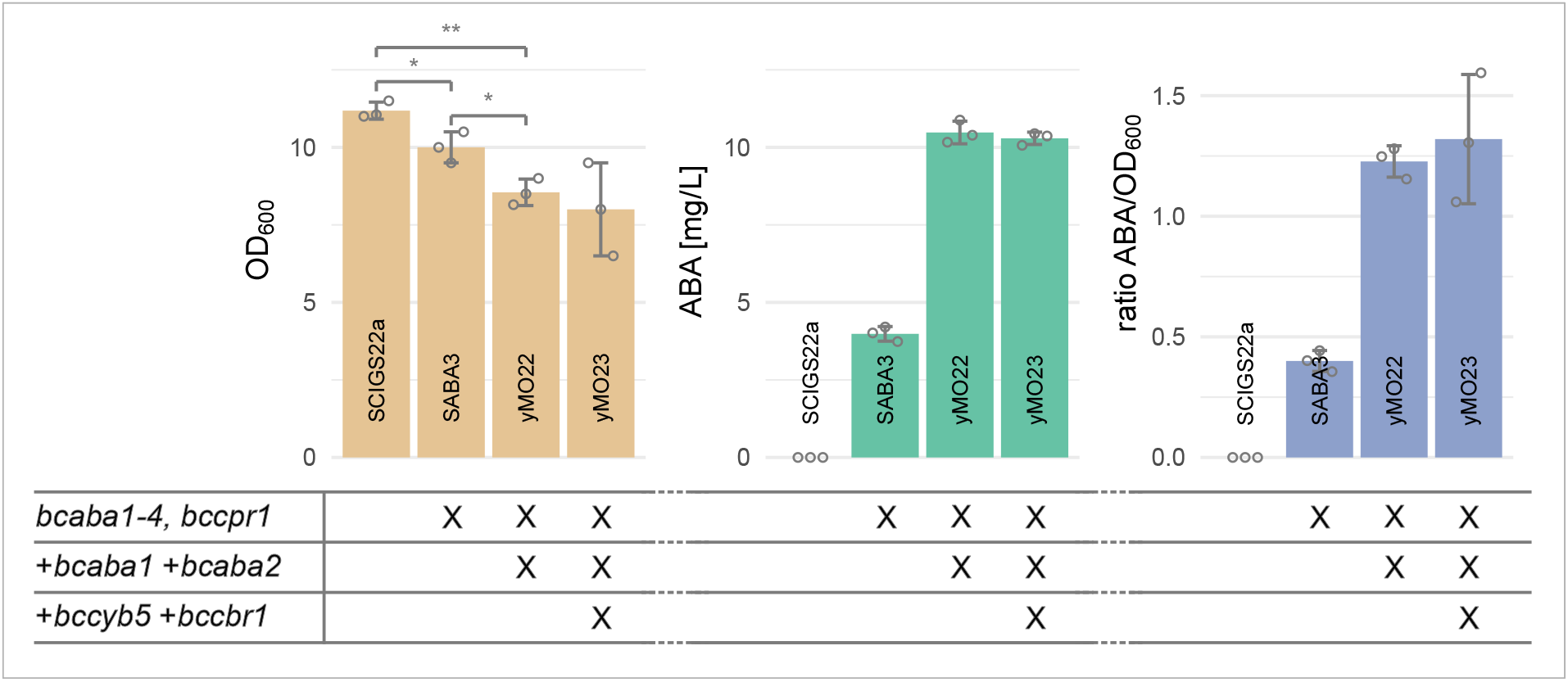
Effects of expressing additional copies of *bcaba1* and *bcaba2*, as well as expressing the *B. cinerea* CYB5 (*bccyb5*) and its cognate CBR (*bccbr1*). “X “ indicates presence of the genetic modification in the strain. OD_600_ (beige), ABA titer in the supernatant (green) and ABA titer normalized to OD_600_ (blue) are shown after 48 h of cultivation in mineral media (24-deepwell microplates). Grey circles show the individual data points used to calculate the mean and standard deviation. No ABA was detected for strain SCIGS22a. When not apparent, significance (Student ‘s t-test, two-sided) is displayed as “* “ and “** ‘ ‘, indicating p<0.05 and p<0.01 respectively.

**Figure 5:**
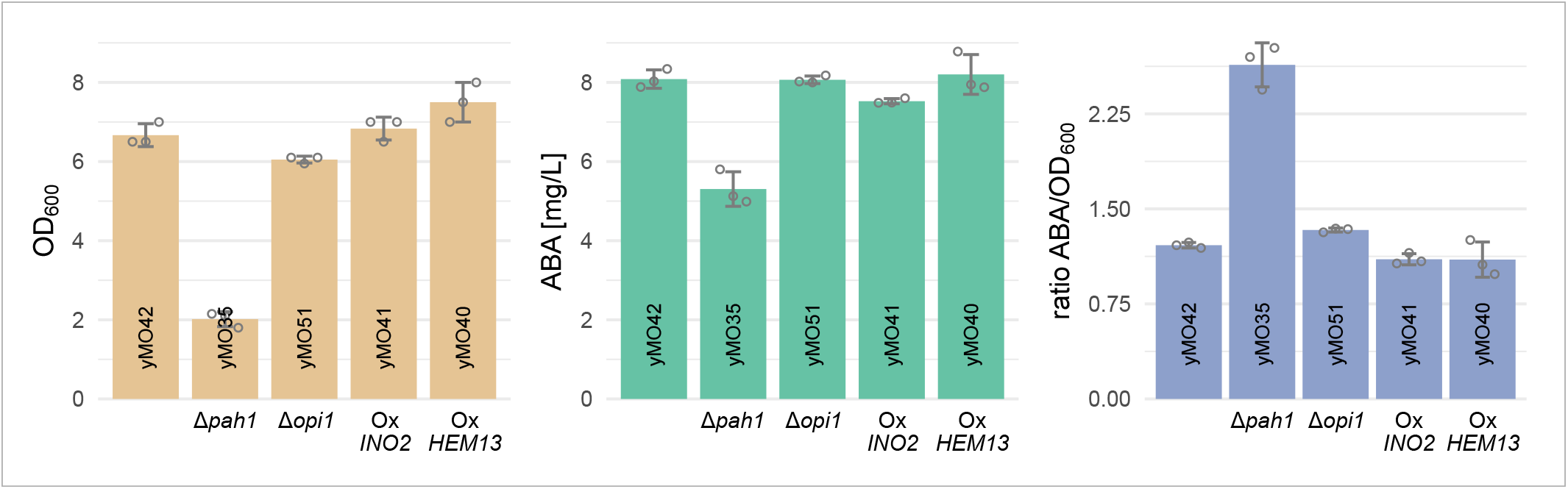
Effects of Δ*pah1*, Δ*opi1, INO2* overexpression and *HEM13* overexpression. Genetic modifications additional to yMO42 (yMO23 carrying an empty plasmid) are displayed below the bars and include knockout of *PAH1* (yMO35) or *OPI1* (yMO51), and episomal overexpression of *INO2* (yMO41) or *HEM13* (yMO40). OD_600_ (beige), ABA titer in the supernatant (green) and ABA titer normalized to OD_600_ (blue) are shown after 60 h of cultivation in mineral media (24-deepwell microplates). Grey circles show the individual data points used to calculate the mean and standard deviation. Ox = overexpression

**Figure 6:**
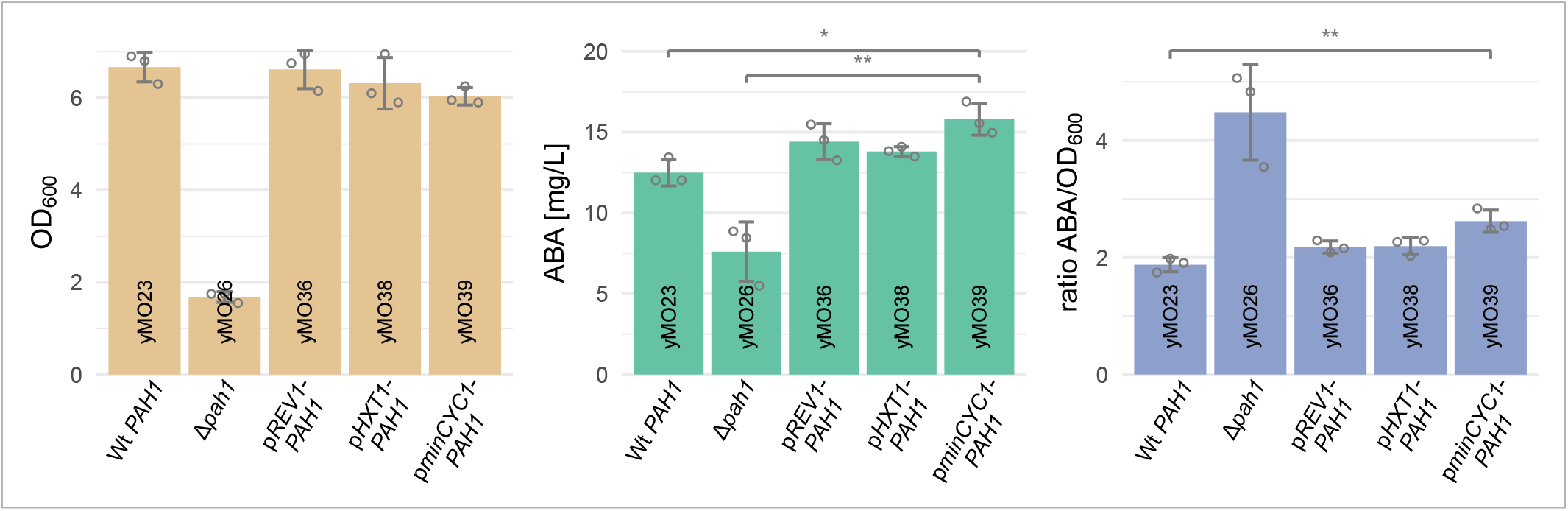
Effects of *PAH1* promoter replacement. The *PAH1* promoter was exchanged for p*REV1* (yMO36), p*HXT1* (yMO38) or p*minCYC1* (yMO39) and the effect compared to the parent strain with wild-type p*PAH1* (yMO23) or a Δ*pah1* strain (yMO26). OD_600_ (beige), ABA titer in the supernatant (green) and ABA titer normalized to OD_600_ (blue) are shown after 60 h of cultivation in mineral media (24-deepwell microplates). Grey circles show the individual data points used to calculate the mean and standard deviation. When not apparent, significance (Student ‘s t-test, two-sided) is displayed as “* “ and “** ‘ ‘, indicating p<0.05 and p<0.01 respectively. Wt = wild-type

### ABA extraction and quantification

ABA was extracted similar to the procedure described in Otto *et al*. 2019. 1 mL of ethyl acetate (>99.9% Sigma-Aldrich) and 0.5% (v/v) formic acid (>98%, Sigma-Aldrich) was added to the 2-mL Eppendorf tubes containing 1 mL culture supernatant. The tubes were vortexed for 10 s, then centrifuged (10 min, 13.000 xg, 4 °C) and 0.8 mL of the supernatant was transferred to a new Eppendorf tube. The solvent was evaporated using a Genevac miVac (45 min, 20 mBar, 45 °C). The pellet was reconstituted in 0.8 mL methanol (>99.9%,

Sigma-Aldrich), centrifuged (10 min, 13.000 xg, 4 °C) and ≈0.5 mL of the supernatant was transferred to an HPLC (high-performance liquid chromatography) vial.

The samples were analysed in an Agilent 6120 Single Quadrupole mass spectrometer (MS) with an Agilent Infinity 1260 HPLC system consisting of a binary pump, autosampler and thermostat column compartment. Ionization was performed using an atmospheric pressure electrospray ionization (API-ES) source (positive mode). Compounds were separated on an Agilent Poroshell 120 EC-C18 (2.7 μm, 3.0 × 50 mm) column (maintained at 40 °C) using a water-acetonitrile gradient with 0.04% formic acid in both solvents. The gradient started with 95% water and, over 5 min, gradually changed to 95% acetonitrile (>99.5%, Sigma-Aldrich). After a 2 min hold, the gradient was ramped back to 95% water over 3 min. The sample injection volume was set to 10 µL. ABA was quantified by selected ion monitoring (m/z = 265) and (S)-(+)-ABA standard (>98%, Cayman Chemicals) was used to fit a calibration curve (2^nd^ polynomial).

### Growth Profiler analysis

Growth (Figure 7) was analysed using a Growth Profiler 960 (Enzyscreen). Precultures were prepared as described above and cultures were subsequently grown in transparent bottom 96-well plate (Enzyscreen) with 250 µL mineral media per well. Pictures were taken every 30 min.

**Figure 7:**
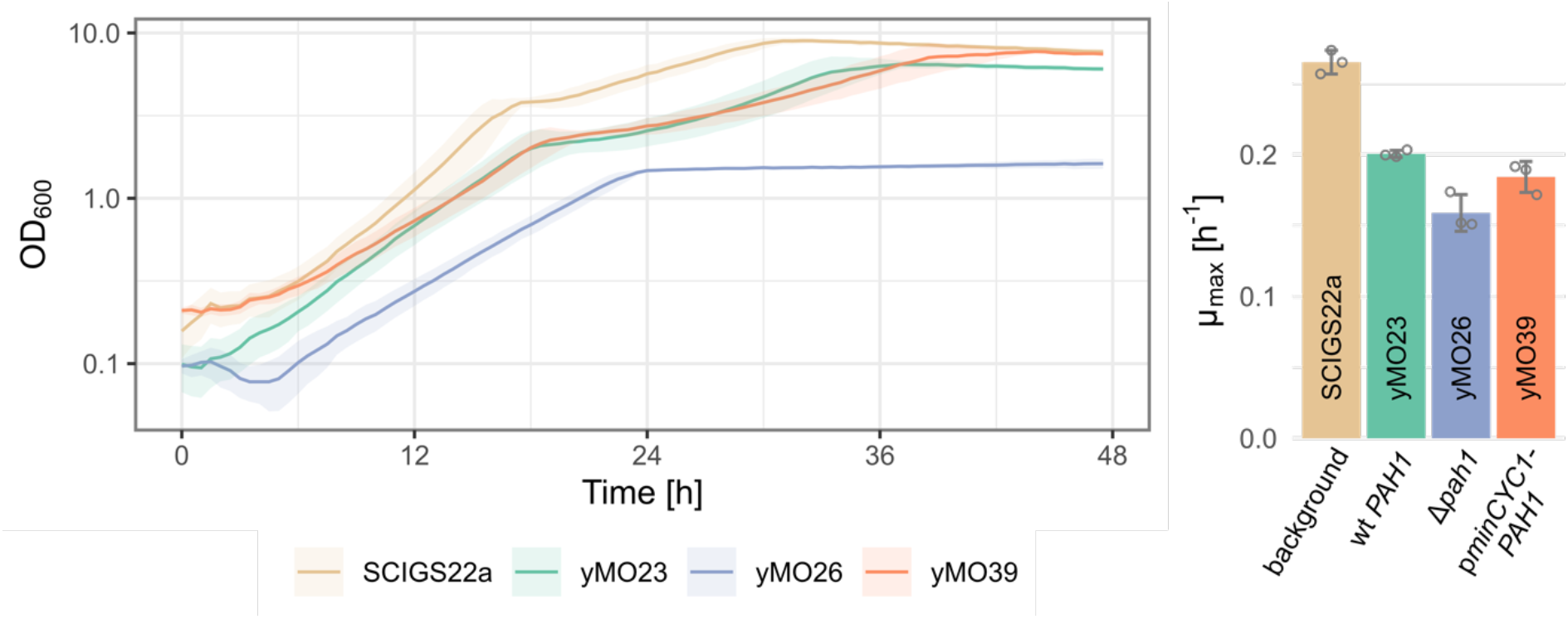
Effect of *PAH1* modifications on cell growth. Growth profiles and maximum growth rate μ_max_ are shown for the background strain SCIGS22a (no *B. cinerea* genes, beige), the wild-type *PAH1* control yMO23 (green), the Δ*pah1* strain yMO26 (blue) and yMO39 carrying the p*minCYC1-PAH1* modification (orange). Lines in the growth profiles show the mean OD_600_ of three replicates and ribbons visualize their standard deviation. Grey circles show the individual data points used to calculate the mean and standard deviation of μ_max_. Cells were grown in mineral media (96-well microplates). Wt = wild-type

### Software

Besides the manufacturer ‘s software for the Growth Profiler (Enzyscreen) and the HPLC-MS (Agilent), R Studio (RStudio Team 2022) was used to analyse the data. Relevant R packages include ggplot2 (Wickham 2016), tidyverse (Wickham *et al*. 2019) and growthrates (Petzoldt 2022). Benchling (www.benchling.com) was used for planning and analysing DNA constructs.

## Results and Discussion

### Effect of transporter overexpression or knock-down on ABA production

Overexpression or knock-down of native transporters could be beneficial for ABA production, either by alleviating product-related cell stress, as was shown for the anti-malaria drug precursor artemisinic acid (Ro *et al*. 2008), or by preventing the export of pathway intermediates.

To test the two hypothesis, two global regulators involved in transporter expression, Yrr1 and Pdr1, were either overexpressed using strong, constitutive promoters, or knocked out. Altered transporter activity in the resulting strains was confirmed by evaluating the response to the antifugal fluconazole. As expected, while overexpression of *PDR1* and *YRR1* resulted in increased fluconazole resistance, their deletion resulted in increased fluconazole sensitivity (Supp. Fig. S1). The role of Pdr1 in the cellular response of yeast to fluconazole has been described previously (Souid *et al*. 2006).

The *B. cinerea* genes *bcaba1, bcaba2, bcaba3, bcaba4* and *bccpr1* were integrated in the strains ‘ genomes to allow ABA production (see Figure 2 for strain pedigree tree). Auxotrophic strains require the transport of essential metabolites from the media into the cell and auxotrophies were shown to greatly affect the cell ‘s transcriptome (Alam *et al*. 2016). To prevent related effects, the analysed strains carried plasmids with auxotrophic marker genes, making them prototrophic. Figure 3 shows the measured OD_600_, ABA titer, ratio of ABA to OD_600_, and ratio of ABA to a compound with the mass-to-charge (m/z) value 233.15, referred to as MZ233. MZ233 was earlier detected in the supernatant of ABA-producing strains and is assumed to be an intermediate or side-product in the ABA pathway (Otto *et al*. 2019). For assessing the activity of transporters, ABA and MZ233 were quantified separately in the supernatant and cell pellet. Small, but significant differences were visible in the measured OD_600_ at 48 h, with yMO49, overexpressing *PDR1* and *YRR1*, growing better compared to the control, yMO48, and the double-knock-out strain, yMO50 (Figure 3). Surprisingly, only traces of ABA and MZ233 were detected in the supernatant and pellet of yMO49. About 50% less ABA was detected in the supernatant for yMO50 compared to the control, however the amount of ABA in the cell pellet remained the same. For the supernatant, the control strain showed a significantly higher ratio of ABA/MZ233 compared to yMO50, though the difference was only 1.2-fold. In contrast, the ABA/MZ233 ratio was 1.7-fold higher for yMO50 in the cell pellet. This indicates that, when normalized to ABA, the double-knock-out strain contains more MZ233 in the supernatant and less MZ233 in the cell pellet.

Neither overexpression of *PDR1* and *YRR1*, nor the double-knockout improved ABA production. In the overexpression strain yMO49, ABA intermediates, likely upstream of MZ233, might be exported rapidly, thereby removing substrates from the product pathway and resulting in only traces being detectable. However, no additional peaks for supposed intermediates were visible in the HPLC-MS chromatogram of the supernatant when comparing yMO49 to yMO48 (Supp. Fig. 2) and high-resolution MS analysis would be necessary to confirm this hypothesis. The increase in OD_600_ for yMO49 indicates that the strain experiences less weak-acid-related stress (Ro *et al*. 2008; Mira, Teixeira and Sá-Correia 2010). The reasons for reduced overall ABA titers in yMO50 compared to the wild-type control are unclear. Nonetheless, assuming that MZ233 is a pathway intermediate, more MZ233 appears to be converted to ABA in yMO50. The results indicate that one or multiple *PDR1*/*YRR1*-regulated transporters facilitate the transport of ABA pathway intermediates if not of the product itself. Pdr1 and Yrr1 control the expression of several ABC transporters involved in the multidrug response in yeast, such as *YOR1, SNQ2, PDR5, PDR10* and *PDR15* (reviewed in Buechel and Pinkett 2020). Modulating the activity of other transcription factors and/or transporters might further impact ABA production. However, a recent study investigated the effect of expressing *Arabidopsis thaliana* ABA transporters in an ABA-producing *Yarrowia lipolytica* strain, aiming to reduce cellular stress, but ABA titers remained unchanged (Arnesen *et al*. 2022).

Taken together, the results suggest that ABA transport is not a promising engineering target to increasing ABA titers in the current strains.

### Genomic integration of additional *bcaba1* and *bcaba2* copies and expression of cytochrome b5

In our previous study, we showed that the CYPs BcABA1 and BcABA2 limit ABA production and that plasmid-based expression of additional gene copies increased the ABA titer more than 4-fold (Otto *et al*. 2019). Genomic integration of expression cassettes is mostly preferred to episomal expression for biotechnological applications, since it results in less cell-to-cell variability and higher genetic stability (Zhang, Moo-Young and Chisti 1996; Lee *et al*. 2015). Therefore, additional copies of *bcaba1* and *bcaba2* were integrated into the genome of SABA3 resulting in the strain yMO22 (see Figure 2 for strain pedigree tree). *Bcaba1* and *bcaba2* were expressed using the strong promoters p*TDH3* and p*CCW12*, respectively. Studies demonstrated that heterologous CYB5 expression can be beneficial for CYP activity (Paddon *et al*. 2013; Ignea *et al*. 2017). We investigated the effect of expressing the *B. cinerea* cytochrome b5, encoded by *bccyb5*, and cognate cytochrome b5 reductase, encoded by *bccbr1*. The strain yMO23 (originating from yMO22) contains an expression cassette with p*HHF2*-*bccyb5* and p*TEF2*-*bccbr1* integrated in the genome.

As was expected, the additional copies of *bcaba1* and *bcaba2* resulted in an increase in ABA production, which was visible in the absolute titer and titer normalised to OD_600_ (Figure 4). The 2.6-fold increase when comparing SABA3 and yMO22 was lower than the 4.1-fold increase that was previously observed in the plasmid-carrying strain (Otto *et al*. 2019). This was anticipated since plasmid gene expression levels are often higher when compared to genomic integrations (Fang *et al*. 2011; Lee *et al*. 2015), likely due to cells carrying on average more than one copy of centromeric plasmids (Singh and Anthony Weil 2002). Expression of *bccyb5* and *bccbr1* in yMO23 did not increase ABA titers. Comparing SABA3 and yMO22, a small but significant decrease in OD_600_ was observable after 48 h of cultivation (Figure 4). Our results indicate that CYP activity was still limiting ABA titers.

However, introducing a third copy of *bcaba1* and *bcaba2* will likely further decrease viability and potentially lead to lower genetic stability. We concluded that modulating the native yeast metabolism would be a more sustainable engineering strategy to improve CYP activity.

### ER proliferation and heme metabolism

ER proliferation is a promising engineering strategy since it was successfully applied for production of other isoprenoids and does not require engineering of the heterologous CYP itself (Arendt *et al*. 2017; Kim *et al*. 2019). Three of the most common native target genes for ER proliferation are *PAH1, INO2* and *OPI1. PAH1* encodes phosphatidate phosphatase, a highly regulated enzyme catalysing the conversion of phosphatidates to diacylgylcerols, a key reaction for balancing levels of membrane phospholipids (PL) and triacylglyceride (TAG) storage lipids (Han, Wu and Carman 2006; Pascual and Carman 2013). Ino2 and Opi1 are transcription factors regulating various lipid metabolism genes. The Ino2/Ino4 complex activates genes involved in PL biosynthesis, whereas Opi1 acts as a repressor when interacting with Ino2 (Henry, Kohlwein and Carman 2012). Knockout of *PAH1* or *OPI1* and overexpression (or deregulation) of *INO2* resulted in similar phenotypes with an enlarged ER (Schuck *et al*. 2009; Arendt *et al*. 2017; Kim *et al*. 2019).

Heme is an essential co-factor for CYPs, CPRs and CYBs. Overexpression of CYPs and their co-enzymes could lead to heme depletion and limit ABA production. A recent study showed that overexpression of the coproporphyrinogen III oxidase gene *HEM13* increases native heme production 3-fold in CEN.PK.113-11C (Ishchuk *et al*. 2022).

We investigated the effects of a *PAH1* or *OPI1* knockout (strains yMO35 and yMO51 respectively) and *INO2* or *HEM13* overexpression (strains yMO41 and yMO40 respectively) (see Figure 2) for strain pedigree tree). Even though the addition of *bccyb5* and *bccbr1* in yMO23 did not result in higher ABA titers (Figure 4), we decided to use the strain for further experiments, since *bccyb5* activity could also be affected by ER expansion and increased heme supply. *INO2* and *HEM13* were overexpressed using centromeric plasmids and the other strains analysed in this experiment were transformed with empty plasmids to render them prototrophic.

Figure 5 shows the OD_600_, ABA titer and ABA titer normalized to OD_600_ for strains with engineered PL metabolism and *HEM13* overexpression. With the exception of the Δ*pah1* strain yMO35, OD_600_ and ABA titers were comparable for all strains. yMO35 exhibited a severe growth defect with ≈70% lower OD_600_ after 60 h compared to the control strain yMO42. Absolute ABA titers for yMO35 were decreased by ≈35%; however, the ABA titer relative to OD_600_ was increased 2.2-fold.

The results indicate that heme supply did not limit CYP activity and that Δ*pah1*-mediated ER expansion was beneficial for ABA production. However, the beneficial effects of the *PAH1* knockout come with impaired growth, a phenotype that has been observed before (Han, Wu and Carman 2006; Zhang *et al*. 2021). Even though modification of the three target genes *PAH1, OPI1* and *INO2* all resulted in ER expansion in previous studies, they did not all affect CYP activity in the ABA-producing strains. A previous study compared the effects of Δ*pah1*, Δ*opi1* and overexpression of *INO2* for the production of isoflavonoids (Liu *et al*. 2021). Flavonoid titers were improved by either knocking out *OPI1* or by overexpressing *INO2*. No beneficial effects were observed for Δ*pah1*, and in this study severe growth defects were only seen for a Δ*pah1*/Δ*opi1* double-knock-out strain and a strain with Δ*opi1* and simultaneous *INO2* overexpression (Liu *et al*. 2021). The beneficial effects of ER proliferation appear to be strain-or product-specific, potentially depending on other genetic modifications or being specific to the heterologous CYP.

The ABA-producing strains of this study are based on the sesquiterpenoid platform strain SCIGS22a, in which the genes *DPP1* and *LPP1* were deleted to prevent the dephosphorylation of farnesyl-pyrophosphate to farnesol (Faulkner *et al*. 1999; Scalcinati *et al*. 2012b; López *et al*. 2015). Like *PAH1, DPP1* and *LPP1* encode phosphatidate phosphatases. Dpp1 and Lpp1 only have minor effects on PL and TAG levels (Carman 2019), but their deletion potentially played a role in influencing the growth and ABA titer of the Δ*pah1* strain yMO35.

### Exchanging of the native *PAH1* promoter to mediate growth deficiency

The deletion of *PAH1* in yMO35 led to a severe growth defect, making the strain unsuitable for further engineering or future biotechnological uses. However, encouraged by the improved relative ABA titer of yMO35 (Figure 5), we investigated if a gene knock-down could mediate the growth defect and still benefit ABA production. Expression levels of the *PAH1* promoter during growth in glucose are comparable to the *REV1* promoter and are <1% of the commonly used strong *TEF1* promoter (unpublished data).

The native *PAH1* promoter was exchanged for three different promoters to investigate their effects on growth and ABA production. We chose the constitutively weak promoter p*REV1* (strain yMO36), the glucose-concentration-dependent promoter p*HXT1* (strain yMO38) and a synthetic minimal promoter named p*minCYC1* (strain yMO39) (Ottoz, Rudolf and Stelling 2014). p*HXT1* is repressed in low glucose conditions and is used in the background strain SCIGS22a to regulate *ERG9* expression with the goal of separating cell growth and production phase (Scalcinati *et al*. 2012a, 2012b). Figure 6 shows the OD_600_, absolute and relative ABA titers of strains with replaced p*PAH1* and control strains with wild-type p*PAH1* (strain yMO23) and Δ*pah1* (strain yMO26). The strains with wild-type *PAH1* and promoter replacement showed no significant difference in OD_600_. The strain carrying the p*minCYC1*-*PAH1* modification, yMO39, produced 15.8 mg/L ABA, ≈1.3-fold the titer produced by the wild-type *PAH1* control yMO23. The ABA titer normalized to OD_600_ for yMO39 was also ≈1.3-fold higher than the titer of yMO23, but remained 40% smaller than for the Δ*pah1* strain yMO26 (Figure 6).

In addition, we compared the growth profile and maximum growth rate μ_max_ of the background strain SCIGS22a, wild-type *PAH1* strain yMO23, the Δ*pah1* strain yMO26 and the p*minCYC1*-*PAH1* strain yMO39 (Figure 7). The growth profiles and μ_max_ of yMO23 and yMO39 were similar, both had a lower μ_max_ and slightly lower final OD_600_ than the background strain. Among the analysed strains, the Δ*pah1* strain yMO26 had the most severe growth defect in terms of μ_max_ and final OD_600_. Interestingly, yMO26 did not show bi-phasic growth. Deletion of *PAH1* has been reported to cause respiratory deficiencies (Irie *et al*. 1993), potentially explaining the growth defect.

In conclusion, by exchanging p*PAH1* for the weak minimal promoter p*minCYC1*, we were able to avoid the growth defect observed in the Δ*pah1* strain yMO26, while still improving ABA titers. The relative ABA titer per OD_600_ was higher for yMO26 than for yMO39, indicating that ABA production might be further improved by fine-tuning ER proliferation.

## Conclusion

Improving CYP activity by protein engineering is challenging and requires in-depth knowledge about the protein and/or high-throughput screening capabilities. In this study, we explored ways to improve heterologous CYP activity by modifying the native yeast metabolism instead of the enzymes themselves.

ER proliferation has proven to be a valuable approach for CYP-expressing cell factories that does not require engineering of the enzymes themselves (Jiang *et al*. 2021). It led to multi-fold increased titers in some cases (Arendt *et al*. 2017; Kim *et al*. 2019). In this study, we showed that the three commonly used target genes for expanding the ER architecture, *PAH1, OPI1* and *INO2*, do not universally benefit ABA production (Figure 5). Only Δ*pah1* lead to an increase in ABA titer when normalized to OD_600_ but caused a severe growth defect. By exchanging p*PAH1* to a weak synthetic promoter, p*minCYC1*, it was possible to mediate the growth defect and still improve ABA titers significantly (Figure 6, Figure 7). While overexpression of *PAH1* had been utilized in TAG cell factories (Hung, Kanehara and Nakamura 2016), to the best of our knowledge, a *PAH1* knockdown had not been investigated before.

Taking the results of the current and previous study (Otto *et al*. 2019) into account, we presume that CYP activity is still a major bottleneck for ABA production. CYP activity might be further improved in the current strain by fine-tuning *PAH1* expression levels. More weak promoters could be tested, or inducible promoters could be used to titrate gene expression, allowing for a systematic investigation of phenotypical effects. Native *PAH1* expression varies depending on the growth phase (Pascual, Soto-Cardalda and Carman 2013) and testing growth-phase specific promoters, like p*HXT1*, in different cultivation conditions, e.g. fed-batch cultivation, would provide valuable insight for biotechnological applications. Another engineering strategy would be combining the *PAH1* knockdown with *INO2* overexpression or *OPI1* deletion.

In a recent study, *Y. lipolytica* was used as a heterologous production host for ABA (Arnesen *et al*. 2022). Complex media or mineral media containing 80 g/L glucose was used, making the titers not comparable to this study. Similar to the Δ*pah1* phenotype in *S. cerevisiae*, deletion of *PAH1* in *Y. lipolytica* results in massive ER expansion (Guerfal *et al*. 2013). We assume that *PAH1* deletion or p*PAH1* replacement could be a promising engineering strategy in a *Y. lipolytica* ABA cell factory.

Few biotechnological studies have investigated the various target genes for ER proliferation comprehensively. This report provides useful data for future studies and is a step towards a more systematic understanding of ER proliferation for improving CYP activity.

## Acknowledgments

We would like to thank the Chalmers Mass Spectrometry Infrastructure (CMSI) for their help and technical support with the HPLC–MS analysis.

## Contributions

MO, FD and VS conceived this project. MO, MG, FD and VS designed the research. MG performed and analysed the fluconazole experiment. MO performed and analysed the other experiments. MO, MG, FD and VS prepared the manuscript.

