## Supplementary Material for "Engineering Yeast to Improve Heterologous Abscisic Acid Production"

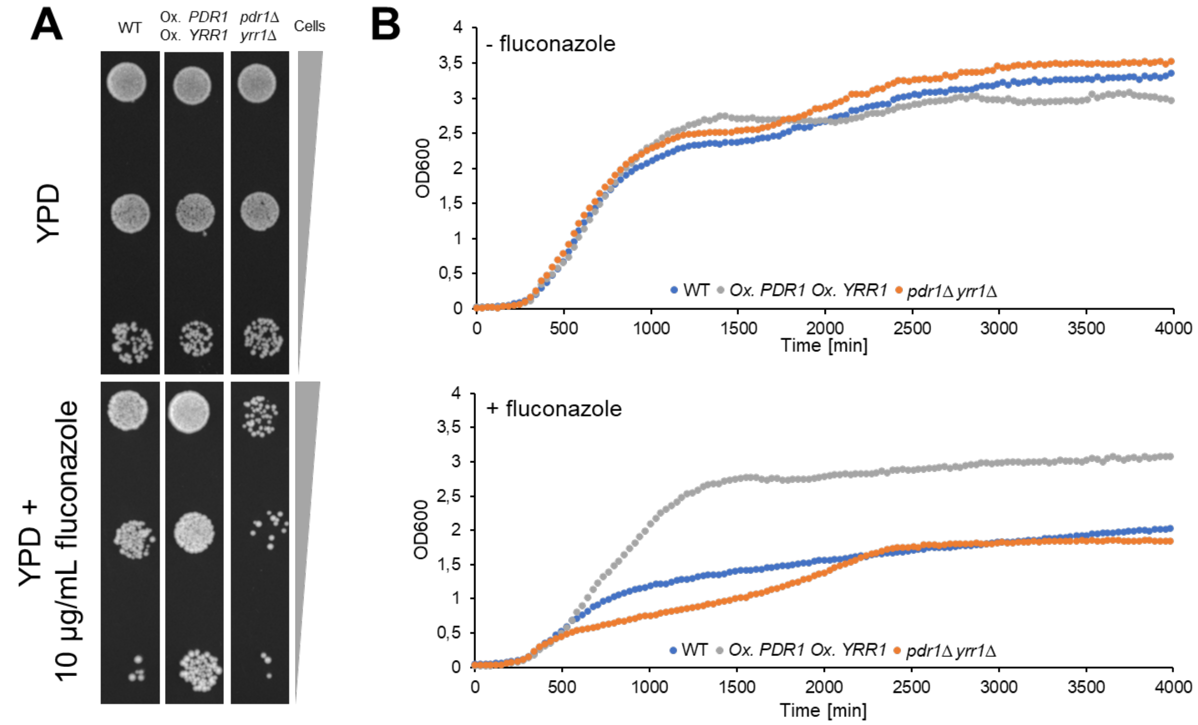


Figure S1: Effect of fluconazole on *PDR1+YRR1* overexpression or knock-out strains. **A** Cells were grown to mid *log* phase (OD≈0.5) in YPD, and diluted to OD=0.05, OD=0.01 and OD=0.001. 5 µL of these dilutions were spotted onto YPD and YPD+10 µg/mL fluconazole agar plates and were cultivated for 3 d at 30 °C. WT, CEN.PK113-11C; Ox. *PDR1* Ox. *YRR1*; yMG01, *pdr1Δ yrr1Δ*, yMG02. **B** Cells were cultivated in 250 µL SD (top panel) and SD+15 µg/mL fluconazole (bottom panel) at 30 °C, in a 96-well microtiter plate. Starting OD=0.01. Growth was monitored in an Enzyscreen Growth Profiler 960. Data points are the mean of technical triplicates.

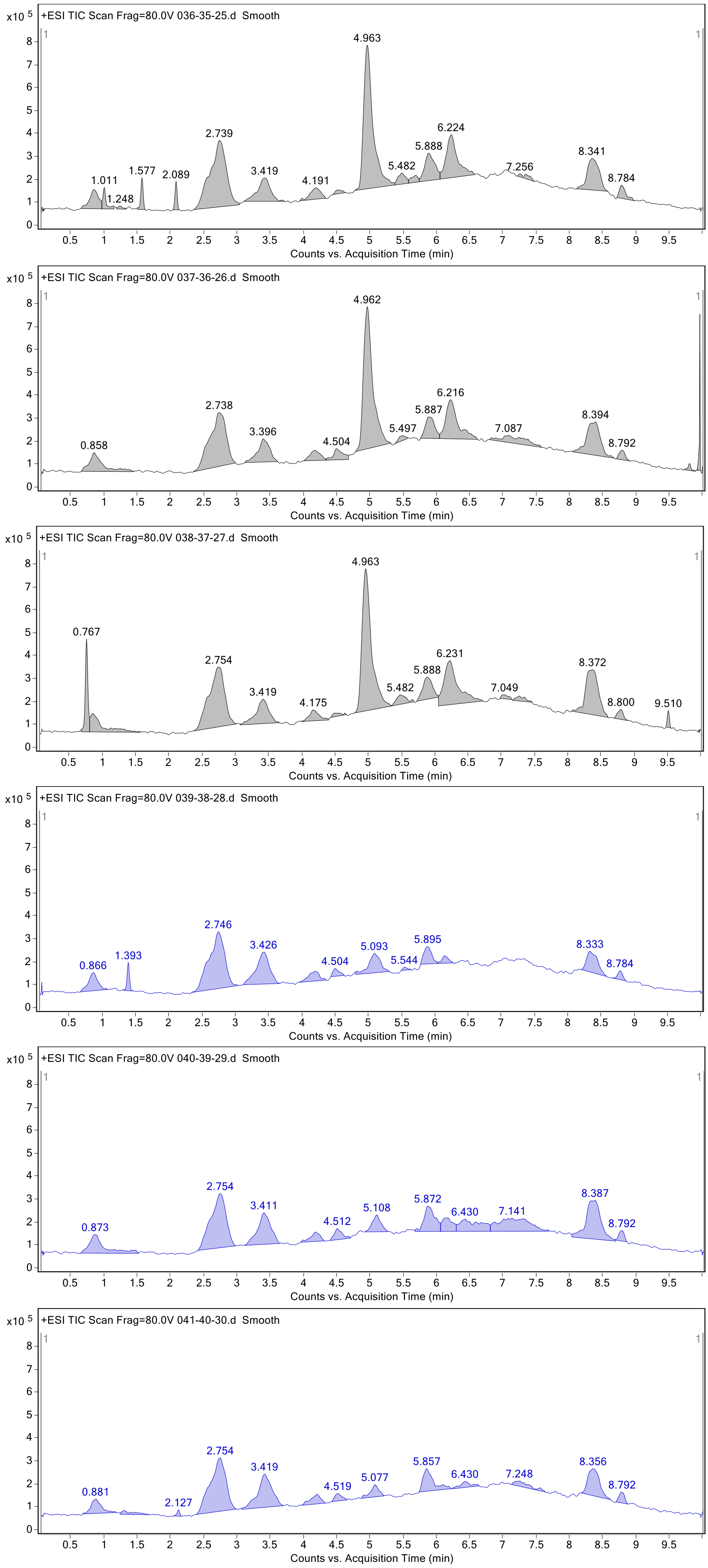


Figure S2: Total ion chromatograms of yMO48 (grey) and yMO49 (blue), 3 replicates are shown per strain

Table S1: Primers used in this study

| Primer name | Sequence |
|---|---|
| 306 | GCATCGTCTCATCGGTCTCATATGTCTAACTCAATCTTGAATTTGGG |
| 307 | ATGCCGTCTCAGGTCTCAGGATCCTTATTTGTATTCTGTACCTTCAG |
| 308 | GCATCGTCTCATCGGTCTCATATGTTGTTATCTATTAAAGATTTG |
| 309 | ATGCCGTCTCAGGTCTCAGGATCCTTATCTTGGAACTTCTTTTAAC |
| 310 | TATCAGAATGAGGCCACCATTGTTGGGT |
| 357 | ACCCAACAATGGTGGCCTCATTCTGATA |
| 366 | GAAGAGTAGGCCATCGTGGG |
| 367 | CAGCAAGCGAGTTTCGTGTG |
| 417 | GGACAAGTGTTCAGAGCAGTATAATGTTGCTTCTGTATCAAATGACCCTAaccgtagtcggtctcaaacg |
| 418 | GAAGACCATGTTTTAGACACAGACCCAAGAGCTCTGCCTACGTACTGCATagatctcgctggatatgcc |
| 422 | GGACAAGTGTTCAGAGCAGTATAATGTTGCTTCTGTATCAAATGACCCTAGCAGGTCCCATCTGGAATATAATTCC |
| 423 | GAAGACCATGTTTTAGACACAGACCCAAGAGCTCTGCCTACGTACTGCATGATTTTACGTATATCAACTAGTTGACGATTATG |
| 424 | GGACAAGTGTTCAGAGCAGTATAATGTTGCTTCTGTATCAAATGACCCTActagagcatgtgctctgtatgtata |
| 425 | GAAGACCATGTTTTAGACACAGACCCAAGAGCTCTGCCTACGTACTGCATaagcttgatatcgaattcctgc |
| 426 | CCAACTCGATTTCGCTCAGC |
| 427 | GCTCTGCCTACGTACTGCAT |
| 1260 | TCCACCTCAATGAGAGGGATTAA |
| 1261 | GGTCCTTACTGAGCCATTCAAAGG |

Table S2: Standard PCR reactions with annealing temperature $T_{ann}$ and elongation time $t_{elo}$ (in min:sec)

| Primers | Template | Polymerase | $T_{ann}$ | $t_{elo}$ | Description |
|---|---|---|---|---|---|
| 292/293 | *bccyb5* (Genscript) | Phusion | 53°C | 0:30 | *bccyb5* attach MoClo overhangs |
| 294/295 | *bccbr1* (Genscript) | Phusion | 53°C | 0:30 | *bccbr1* attach MoClo overhangs |
| 306/357 | p416-ABA1 (Otto *et al.* 2019) | PrimeStar | 55°C | 1:15 | *bcaba1* remove BsaI site and attach MoClo overhangs |
| 307/310 | p416-ABA1 (Otto *et al.* 2019) | PrimeStar | 55°C | 1:15 | *bcaba1* remove BsaI site and attach overhangs |
| 308/309 | p416-ABA2 (Otto *et al.* 2019) | PrimeStar | 55°C | 2:15 | *bcaba2* attach MoClo overhangs |
| 366/367 | *S. cerevisiae* genomic DNA | SapphireAmp | 55°C | 0:35 | validation Δ*pah1* |
| 417/418 | pYTK027 (Lee *et al.* 2015) | PrimeStar | 55°C | 1:30 | repair cassette p*REV1* |
| 422/423 | *S. cerevisiae* genomic DNA | PrimeStar | 55°C | 1:30 | repair cassette p*HXT1* |
| 424/425 | FRP793_insul-(lexA-box)4-PminCYC1-Citrine-TCYC1 (Ottoz, Rudolf and Stelling 2014) | PrimeStar | 55°C | 0:20 | repair cassette p*minCYC1* |
| 426/427 | *S. cerevisiae* genomic DNA | SapphireAmp | 55°C | 0:20 | validation p*PAH1* |
| 1260/1261 | *S. cerevisiae* genomic DNA | SapphireAmp | 50°C | 0:20 | validation Δ*opi1* |

Table S3: List of MoClo assemblies. Protocol according to Lee *et al.* (2015) and Otto *et al.* (2021)

| Plasmid name | MoClo parts used | MoClo level | Note |
|---|---|---|---|
| pMC3-bcaba1 | pYTK001<br>PCR product 306/307 | 0 | |
| pMC3-bcaba2 | pYTK001<br>PCR product 308/309 | 0 | |
| pMC3-bccyb5 | pYTK001<br>PCR product 292/293 | 0 | |
| pMC3-bccbr1 | pYTK001<br>PCR product 294/295 | 0 | |
| pMC-Ura-Cen | pYTK002<br>pYTK047<br>pYTK067<br>pYTK074<br>pYTK081<br>pYTK081 | 1 (backbone) | contains GFP dropout |
| pMC-His-Cen | pYTK003<br>pYTK047<br>pYTK072<br>pYTK076<br>pYTK081<br>pYTK083 | 1 (backbone) | contains GFP dropout |
| pMMC16 | pMC-Ura-Cen<br>pYTK009<br>pMC3-bcaba1<br>pYTK056 | 1 | |
| pMMC17 | pMC-His single<br>pYTK010<br>pMC3-bcaba2<br>pYTK055 | 1 | |
| pMMC24 | pMC-Ura single<br>pYTk12<br>pMC3-bccyb5<br>pYTK054 | 1 | |
| pMMC25 | pMC-His single<br>pYTK14<br>pMC3-bccbr1<br>pYTK052 | 1 | |
| pX3-bcaba1+2 | pMC-X3<br>pMMC16<br>pMMC17 | 2 | |
| pXII2-bccyb5+cbr1 | pMC-XII2<br>pMMC24<br>pMMC25 | 2 | |

Table S4: Media composition

| | |
|---|---|
| **YPD** | |
| yeast extract (Merck) | 10 g/L |
| peptone from meat (Merck) | 20 g/L |
| glucose (Merck) | 20 g/L |
| for plates: agar agar (Merck) | 20 g/L |
| **SD (pH adjusted to 6 with KOH)** | |
| complete supplement mix dropout -uracil (Formedium) | 0.77 g/L |
| yeast-nitrogen base without amino acids (Formedium) | 6.9 g/L |
| glucose (Merck) | 20 g/L |
| for plates: agar agar (Merck) | 20 g/L |
| **LB (pH adjusted to 6.5 with NaOH)** | |
| peptone from casein (Merck) | 10 g/L |
| NaCl (Merck) | 10 g/L |
| yeast extract (Merck) | 5 g/L |
| for plates: agar agar (Merck) | 20 g/L |
| **Mineral media (pH adjusted to 6.5 with KOH)** | |
| ammonium sulfate (Merck) | 7.5 g/L |
| monopotassium phsophate (Merck) | 14.4 g/L |
| magnesium sulfate heptahydrate (Merck) | 0.5 g/L |
| glucose (Merck) | 20 g/L |
| trace metal solution | 2 mL/L |
| vitamin solution | 1 mL/L |
| for plates: agar agar (Merck) | 20 g/L |
| for auxotrophic strains: uracil (Alfa Aesar) | 100 mg/L |
| **Trace metal solution** | |
| $FeSO_4 \bullet 7H_2O$ | 3 g/L |
| $ZnSO_4 \bullet 7H_2O$ | 4.5 g/L |
| $CaCl_2 \bullet 2H_2O$ | 4.5 g/L |
| $MnCl_2 \bullet 4H_2O$ | 1 g/L |
| $CoCl_2 \bullet 6H_2O$ | 300 mg/L |
| $CuSO_4 \bullet 5H_2O$ | 300 mg/L |
| $Na_2MoO_4 \bullet 2H_2O$ | 400 mg/L |
| $H_3BO_3$ | 1 g/L |
| KI | 100 mg/L |
| $Na_2EDTA \bullet 2H_20$ | 19 g/L |
| **Vitamin solution** | |
| D-Biotin | 50 mg/L |
| D-Pantothenic acid hemicalcium salt | 1 g/L |
| thiamin-HCl | 1 g/L |
| pyridoxin-HCl | 1 g/L |
| nicotinic acid | 1 g/L |
| 4-aminobenzoic acid | 0.2 g/L |
| myo-Inositol | 25 g/L |

Table S5: Oligonucleotides used in this study. gRNA target sequences are underlined.

| Oligo description | Sequence |
|---|---|
| Repair oligo Δ*pah1* | TTTTACCTTCTAAGAAACATACAGGGAAGACATTACTGAAGATAGACACATCGGTCGATTAGATTCTTGTAGCCGAATATTATTTATAACGATCCATACTGCATATTAAGTAAAATTATG |
| gRNA fragment Δ*pah1* | TGCGCATGTTTCGGCGTTCGAAACTTCTCCGCAGTGAAAGATAAATGATCTGATTGAGGGGGGCTTGATGGTTTTAGAGCTAGAAATAGCAAGTTAAAATAAGGCTAGTCCGTTATCAAC |
| Repair oligo Δ*opi1* | TTAAAGCGTGTGTATCAGGACAGTGTTTTTAACGAAGATACTAGTCATTGCCTCTAATACATCCAACACTCTACGCCCTCTTCAAGAGCTAGAAGGGCACCCTGCAGTTGGAAAGGGAATTATTTCGTAAGGCGAGCCCATACCGTCATTCATGCGGAAGAGTTAACACGATTGGAAGTAGGAATAGTTTCGAACCACGGTTACTAATCCTAATAACGGAACGCTGTCTGAAGGATGAGTGTCAGCGAGTGTAACTCGATGAGCTACCCAGTAGTCGTACTGGTCGAGACAACATTGTACCCCAGCGGCGGCGCGGCCAGCTCTAATGCACTCAATCCCGAGGCCTGACGCGACATATCAGCTTAGACTAGGGCGGGGGTGTTGACGTTTGGGGTTGAATAAATCTATTGTACTAATCGGCTTCAACGTGCCCCACGGGTGGCACCTCAGGAGGGGCCCACAGCGAGGAAGTAAACTGTTATTCGTCGGCGATGGTGGTAGCTAATTATGTTCCTTGCCACTACAATAGTATCTAAGCCGTGTAATGGGAACATCCACACCCGAGACAGATTGAGGTCTTTCATGCATTACCACCAGTAATAATATTATA |
| gRNA fragment Δ*opi1* | TGCGCATGTTTCGGCGTTCGAAACTTCTCCGCAGTGAAAGATAAATGATCTGTCGCGGGCGATTGCCAAGGTTTTAGAGCTAGAAATAGCAAGTTAAAATAAGGCTAGTCCGTTATCAAC |
| gRNA fragment p*PAH1* replacement | TGCGCATGTTTCGGCGTTCGAAACTTCTCCGCAGTGAAAGATAAATGATCTTAGAGAATGAGCAGCACGTGTTTTAGAGCTAGAAATAGCAAGTTAAAATAAGGCTAGTCCGTTATCAAC |
